# Cell walls are dynamically O-acetylated in the green seaweed, Ulva compressa

**DOI:** 10.1101/2022.05.24.493306

**Authors:** John H. Bothwell, Alexander J. Goodridge, Marie Rapin, Patrick J. Brennan, Alexandra Traslaviña López, Akanksha Agrawal, Stephen C. Fry, Georgia Campbell, Jonas Blomme

## Abstract

We report that cell wall polymers in the chlorophyte algae may be modified by *O*-acetylation. The importance of cell wall modifications in the green algae is not well understood, although similar modifications play key roles in land plants by modulating the properties of cell wall carbohydrate polymers. Using a combination of biophysical (Fourier-transform infrared and cross-polarisation heteronuclear correlation nuclear magnetic resonance spectroscopy), biochemical (thin-layer chromatography) and molecular approaches (yellow fluorescent protein-tagged transgene localisation), we show that the extractable ulvan fractions of *Ulva compressa* cell walls contain *O*-acetyl sidechains, we demonstrate that acetylation is dynamic and decreases reversibly in response to metal-induced stress, we note interactions between acetyl and borate sidechains and we locate two candidate genes that, together, may encode an acetyltransferase. We therefore propose that *O*-acetylation of ulvan residues is involved in the normal cell wall physiology of at least some chlorophyte algae. To the best of our knowledge, this is the first demonstration of *O*-acetyl sidechains in green algal cell wall polymers, and of reversible changes in algal cell wall polymer modification in response to stress.

## Introduction

*O*-Acetylation is the substitution of an acetyl group (CH_3_CO-) for the hydrogen of an alcohol group (ROH) to form an acetic ester (CH_3_COOR). It is a common but relatively underexplored cell wall modification, in which acetyl groups may be ester linked to -OH groups on sugars or lignins (for reviews, see Gille and Pauly, 2012; Pauly and Ramírez, 2018). Cell wall *O*-acetylation has been reported across several kingdoms, from bacteria (Franklin et al., 2011) to fungi (Levine et al., 1959) to land plants (York et al., 1984).

As with most cell wall modifications, the role of *O*-acetylation remains opaque, but it is often associated with changes in cell wall function (Pauly and Ramírez, 2018). Acetyl sidechains may act by blocking the addition of other substituents such as xylosyl groups (Liu et al., 2016), which would be comparable to the way in which they inhibit protein kinases by blocking phosphorylation sites (Mukherjee et al., 2006), or they may simply alter the bulk rheological properties of cell wall polymers (Schmelter et al., 2002). The extent to which these two mechanisms contribute to the effects of *O*-acetylation remains unclear and may vary from case to case.

Nevertheless, despite any uncertainty over the function of this modification, significant progress has been made in elucidating the molecular machinery responsible for cell wall *O*-acetylation. The extent of acetylation is thought to reflect a balance between acetyl addition by Cas1p-like *O*-acetyltransferases in the golgi lumen (Pauly and Scheller, 2000; Obel et al., 2009) and acetyl removal by acetylesterases in the apoplast (Zhang et al., 2017). The Cas1p protein was discovered in the fungus, *Cryptococcus neoformans* (Janbon et al., 2001), and the only known function of members of this gene family is the attachment of acetyl sidechains to sugar residues (Lee et al., 2011). Its fungal, mammalian and bacterial members are predicted to contain two main domains: first, a series of 11-12 transmembrane helices that are presumed to transport acetyl groups from the cytosol into the golgi lumen and, second, a lumen-facing loop, which contains characteristic GDS and DxxH motifs that are thought to add acetyl groups to their targets (Gille and Pauly, 2012).

In land plants this machinery is slightly more complex because the two domains of Cas1p have been separated into two distinct proteins. The first, transmembrane, domain finds land plant orthologs in the Reduced Wall Acetylation (*RWA*) family, which has four members in Arabidopsis (Manabe et al., 2011). The second, lumen-facing loop, domain is found in the single Arabidopsis Altered Xyloglucan 9 (*AXY9*) gene (Schultink et al., 2015) and in the closely related Trichome Birefringence-Like (*TBL*) gene family, which contains around 50 members in Arabidopsis (Pauly and Ramírez, 2018). Because of the importance of the cell wall to land plants, it is generally argued that the diversity of Trichome Birefringence-Like genes reflects some degree of substrate specificity, with specific genes being responsible for the acetylation of specific cell wall polymers (Gille et al., 2011; Pauly and Ramírez, 2018).

Outside of land plants, the role of cell wall *O*-acetylation in the green lineage is less well established (Pauly and Ramírez, 2018). Published analyses of chlorophyte cell wall fractions do not show any obvious signals from acetic esters (for example Trivedi *et al*., 2016) and while the genome of the model chlorophyte *Chlamydomonas* possesses a Reduced Wall Acetylation-family ortholog, it lacks any obvious *AXY9* or Trichome Birefringence-Like orthologs (Pauly and Ramírez, 2018).

It is, however, unfair to expect *Chlamydomonas* to act as a model organism for the study of cell walls across all chlorophyte algae because its cell wall is constructed from glycoproteins (Roberts et al., 1972; Catt et al., 1976), while those of the green chlorophyte seaweeds are constructed from polysaccharides (Percival, 1979; Baudelet et al., 2017). There is also considerable diversity within the polysaccharide walls of the green seaweeds: the Ulvales, such as the recently sequenced *Ulva compressa* (De Clerck et al., 2018) and *Ulva prolifera* (He et al., 2021), have plant-like cell walls in which cellulose is combined with hemicelluloses and the pectin-like ulvan (Percival, 1979). The Bryopsidales, on the other hand, represented by the recently-sequenced *Caulerpa lentillifera* (Arimoto et al., 2019), have cell walls in which (1→3)-xylan is the main microfibrillar polysaccharide (Frei and Preston, 1964), with no detectable cellulose and no obvious orthologs of cellulose synthase.

Accordingly, in this study we investigate the model green seaweed, *Ulva compressa*, and report evidence for the *O*-acetylation of its extractable cell wall fractions. We show that this acetylation may change reversibly in response to imposed metal stress, identify *Ulva* orthologs to the Arabidopsis *RWA* and *AXY9* acetyltransferase genes and use YFP-tagged transgenic lines to show that these *Ulva* orthologs are localised to the golgi. Gene knockout techniques have not yet been established in *Ulva* but we demonstrate that a strong cell wall acetylation signal and the putative acetyltransferase genes are both present in the ulvalean *U. compressa* and both absent in the bryopsidalean *Caulerpa lentillifera*. Taken together, our results suggest that *O*-acetylation plays a previously unremarked role in the physiology of ulvophyte cell walls.

## Results

### A land plant cell wall fractionation protocol gives reliable separation of chlorophyte polymers

Cell wall fractionation protocols are well established in land plants but have been less well developed for the chlorophyte algae. *Ulva* cell walls contain non-extractable cellulose and a range of extractable polysaccharides whose exact composition varies between species (Percival, 1979). Their major extractable polysaccharide, ulvan, is a heterogeneous polymer built primarily from five sugar monomers: L-rhamnose (Plant and Johnson, 1941), D-glucose, D-xylose, D-glucuronic acid (all Brading, Georg-Plant and Hardy, 1954) and L-iduronic acid (Quemener, Lahaye and Bobin-Dubigeon, 1997). *Ulva* cell walls also contain significant amounts of alkali-extractable phycoxyloglucans and smaller amounts of polymers that yield arabinose, galactose, and mannose, which are presumed to be arabinogalactan- and mannan-like chains (Domozych et al., 2012).

To separate out these polymers, we adapted the protocol of O’Rourke and colleagues (2015) to divide *Ulva* cell walls into several major fractions (Fig. 1). Broadly, these correspond to two fractions that were respectively rapidly and slowly extractable in oxalate buffer and a residue that was not extractable in oxalate (Fig. 1; the three boxes to the top right). The residue that was not extractable in oxalate could be further separated into alkali-extractable and non-extractable fractions (Fig. 1; the four boxes to the bottom right). Thin-layer chromatography verified that extracted fractions contained the expected monosaccharide residues (Fig. 2). The oxalate-extractable fractions had high levels of uronic acids and rhamnose, which are characteristic of ulvan (Percival, 1979). The alkali-extractable fractions contained high levels of xylose, glucose and mannose, which are characteristic of hemicelluloses (Percival, 1979), and the final non-extractable fraction consisted almost entirely of glucose (Fig. 2). Following the naming convention of O’Rourke and colleagues (2015), we will refer to the oxalate-extractable fractions as ulvan 1 and ulvan 2, to the major alkali-extractable fraction as hemicellulose b and to the non-extractable fraction as α-cellulose.

**Figure 1.**
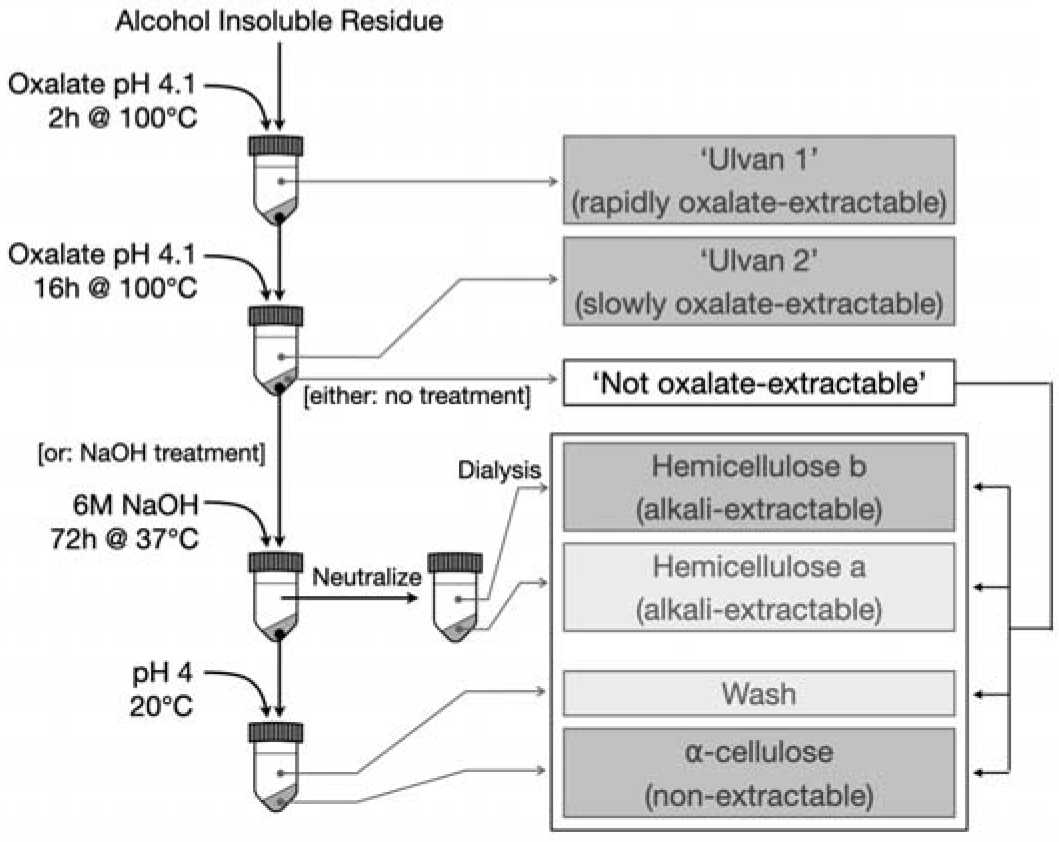
Fractionation of chlorophyte cell walls into broad polymer classes (after O’Rourke *et al*., 2015). The four major extractable fractions are show in darker boxes to the right; the hemicellulose a and Wash fractions were analysed but were found to contain very little material, so are not discussed further. Some investigations used the washed ‘not oxalate-extractable’ residue without subjecting it to alkali hydrolysis (see Supplemental Fig. S7).

**Figure 2.**
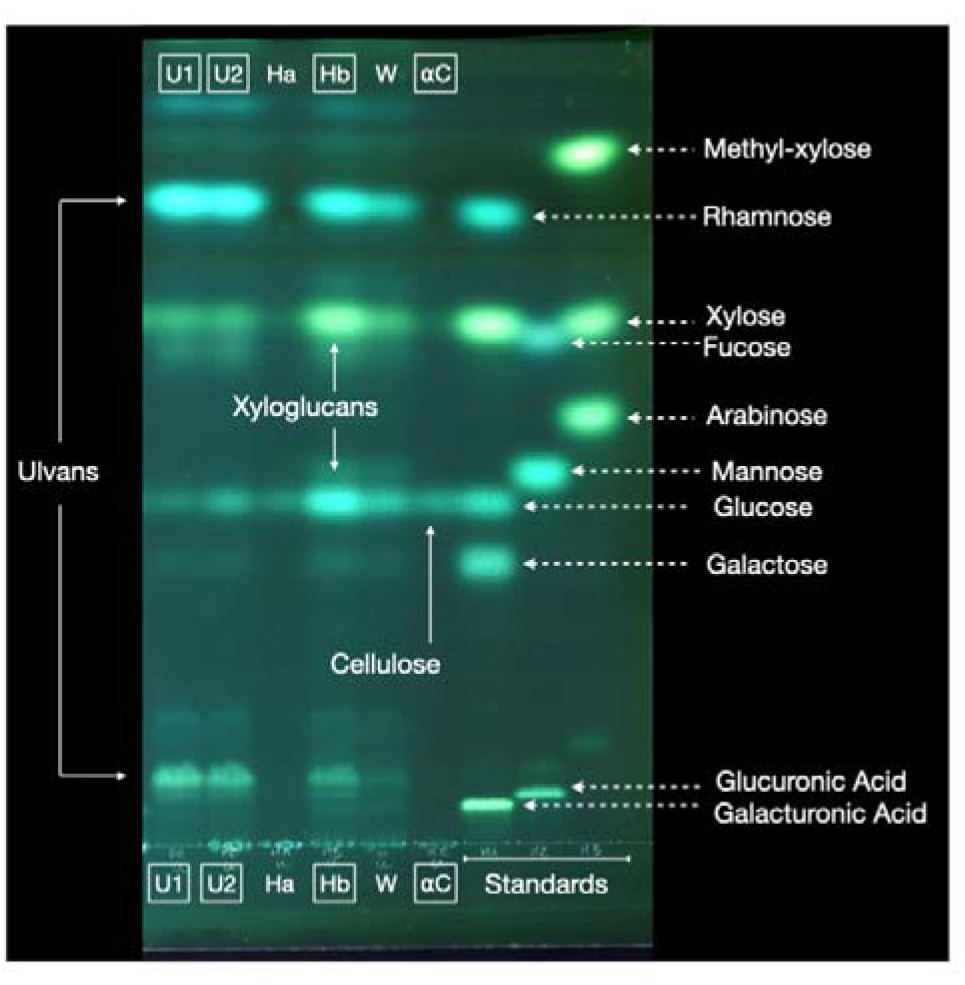
Thin-layer chromatography of the monosaccharide constituents of *Ulva* cell wall fractions. The four major fractions are outlined. Cell wall fractions are ‘ulvans’ (U1, U2), hemicelluloses a and b (Ha, Hb), the wash after alkali extraction (W) and α-cellulose (αC). Chromatography was performed on silica gel in ethyl acetate/pyridine/acetic acid/H2O (6 : 3 : 1 : 1). Stain: thymol/H_2_SO_4_. Monosaccharide standards are shown to the right of the plate and likely parent polymers are indicated with solid arrows. Note that the apparently low α-cellulose yield will be an underestimate because cellulose is not efficiently hydrolysed by the trifluoroacetic acid used for polymer hydrolysis.

### FTIR-visible ester groups are associated with the slowly extractable cell wall fraction in Ulva (‘ulvan 2’)

To look for evidence of *O*-acetylated polymers, we ran FTIR spectra for each cell wall fraction (Fig. 3). The alkaline hydrolysis treatments used in the preparation of the hemicellulose fractions would be expected to remove acetic esters in those fractions, so we also ran spectra for the ‘Not oxalate-extractable’ residue (Fig. 1), which was not subjected to NaOH treatment (Supplemental Fig. S7). The FTIR spectra of polymer mixtures are invariably complex and ours contain multiple overlapping peaks with four prominent clusters centered at ∼1620cm^−1^, ∼1430cm^−1^, ∼1220cm^−1^ and ∼1030cm^−1^ (Fig. 3). The peaks at ∼1620cm^−1^ and ∼1430cm^−1^ contain stretching (ν C=O) and bending (δ COOH) resonances respectively from carboxylic acids (McCann et al., 1994), the peaks ∼1220cm^−1^ contain sulfate stretching resonances (ν S=O; Mainreck *et al*., 2011) and the peaks around ∼1030cm^−1^ contain multiple overlapping resonances from sugar C-O stretches (ν C-O; McCann *et al*., 1994). We tentatively assign the ulvan ∼985cm^−1^ peak to the in-plane rocking from methyl groups (ρ CH_3_; Marry et al., 2000). All of these assignations are supported by our thin-layer chromatograms (Fig. 2) and by the knowledge that, as observed (Fig. 3), more sulfation is expected on the ulvan fraction (Percival and Wold, 1963; Bilan et al., 2007).

**Figure 3.**
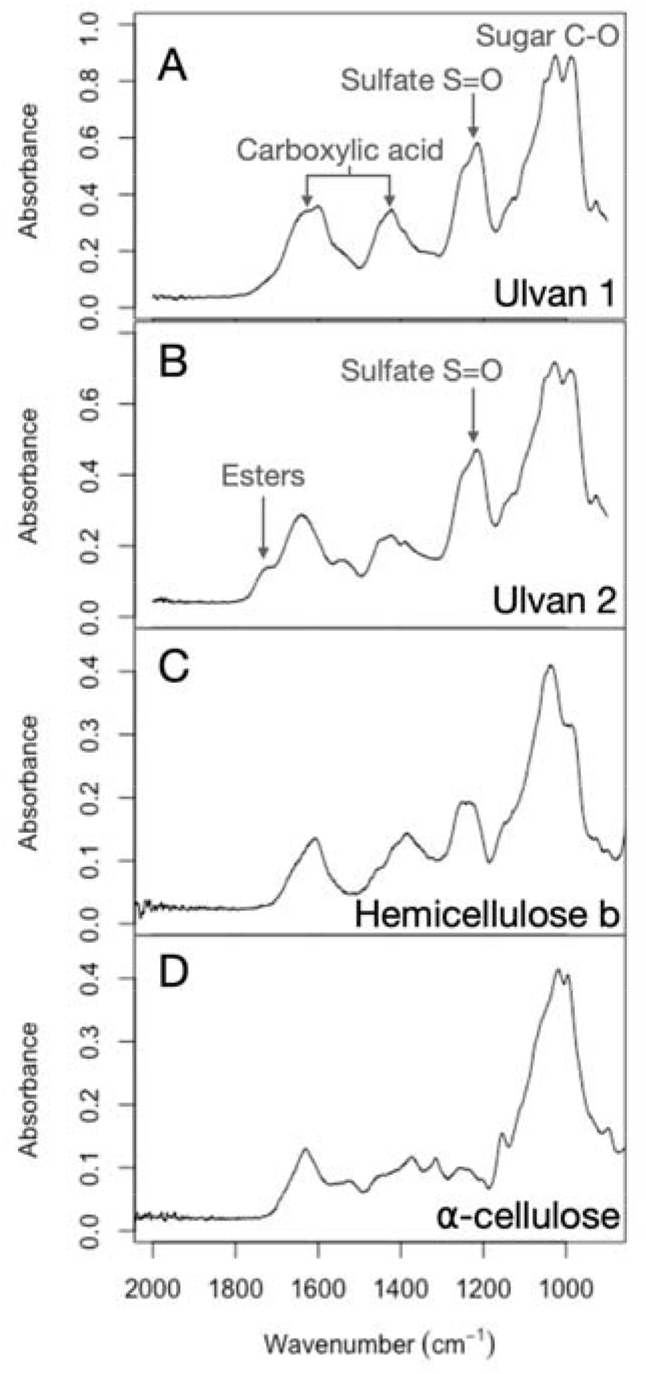
Representative FTIR spectra of *Ulva* cell wall fractions, showing the major resonances between 2000-900 cm^−1^. (A) ulvan 1 fraction, (B) ulvan 2 fraction, (C) hemicellulose b fraction (see also Supplemental Fig. S7) and (D) the α-cellulose fraction. Easily identifiable resonances are indicated with arrows.

Importantly, the slowly oxalate-extractable ulvan 2 faction displays a noticeable additional peak at around ∼1730cm^−1^ (Fig. 3B), which appears as a shoulder on the more prominent carboxylic acid peak cluster centered at ∼1620cm^−1^. Resonances in this region correspond to acetic esters in the pectin fractions of land plant cell walls (McCann et al., 1994) and we give them the same assignment here. We note in passing that *O*-acetylation may contribute to the ulvan 2 fraction’s slower extractability in oxalate buffer than the ulvan 1 fraction because *O*-acetyl groups will form hydrogen bonds with water less readily than the -OH groups that they replace.

### Cross-polarisation heteronuclear correlation (CP-HETCOR) NMR spectra of Ulva cell wall extracts shows magnetization transfer between alkyl and carboxyl groups

To confirm that the ester FTIR signal was due, at least in part, to *O*-acetic esters, we carried out CP-HETCOR NMR spectroscopy of *Ulva* cell wall extracts. These display a variety of crosspeaks in short contact time (0.1ms) CP-HETCOR spectra and, given the sugar residues that are known to be found in *Ulva* cell walls, the majority of crosspeaks may be confidently assigned to particular H-C couplings (Fig. 4A).

**Figure 4.**
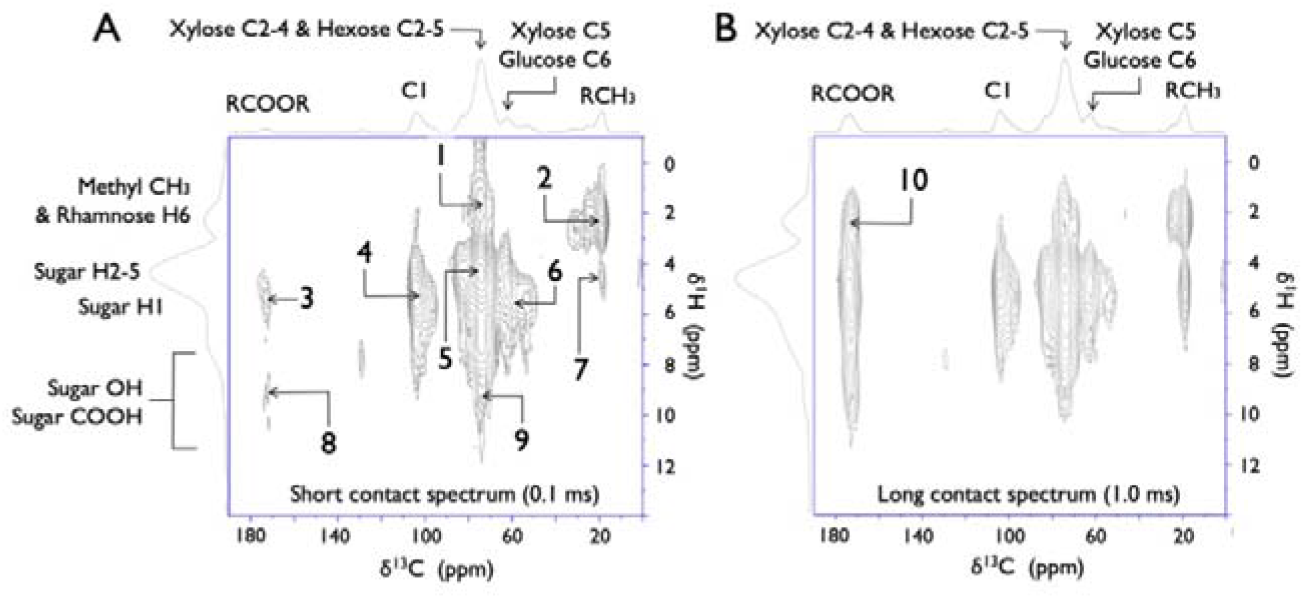
Representative ^13^C CP-HETCOR NMR of *Ulva* cell wall extract, with skyline plots above and to the left of each spectrum. (A) Representative short contact spectrum (0.1ms contact time), (B) Representative long contact spectrum (1.0ms contact time). Crosspeaks indicate magnetization transfer from H atoms to C atoms. Probable H to C transfers are 1: Rha H6 to Rha C5, 2: Rha H6 to Rha C6, 3: Uronic Acid (UroA) H5 to UroA C6, 4: sugar H1 to sugar C1, 5: multiple sugar H2-H5 to sugar C2-C5, 6: Xyl H5 to C5 and Glc H6 to C6, 7: Rha H5 to Rha C6, 8: UroA H6 to UroA C6, 9: multiple sugar OH2-5 to sugar C2-C5; 10: H on CH_3_ to C on COOR. Direct quantification is not trivial with this kind of NMR experiment, but in fully relaxed 1D spectra the carboxylic acid/ester peak at 170-180ppm is consistently comparable in size to the C1 peak at 100-110ppm (cf. skyline plot above Fig. 4B).

When the CP-HETCOR contact time was lengthened from 0.1ms to 1ms, which is long enough to allow two-bond magnetization transfer (Apperley et al., 2012), a prominent new crosspeak appeared (marked as cross-peak #10 to the top left of Fig. 4B): the relevant cross-peak represents magnetization transfer from ^1^H nuclei that resonate between 1.5-2.5ppm (ie protons in alkyl or methyl groups) to ^13^C nuclei that resonate between 170-180ppm (ie carbons in carboxylic acid or alkyl ester groups). The simplest interpretation for this observed cross-polarization is two-bond coupling along an acetyl chain, from a methyl proton through its methyl carbon to a carboxyl carbon (Fig. 5A). We presume that any such acetyl chains reflect the presence of *O*-acetic esters on the C2 and C3 positions of ulvan sugar residues, because there are no alkyl or methyl groups within two bonds of a carboxylic acid group in the unmodified sugar monomers that have been reported in cellulose and ulvan (Figure S1; cf. Lahaye and Robic, 2007; Domozych *et al*., 2012).

**Figure 5.**
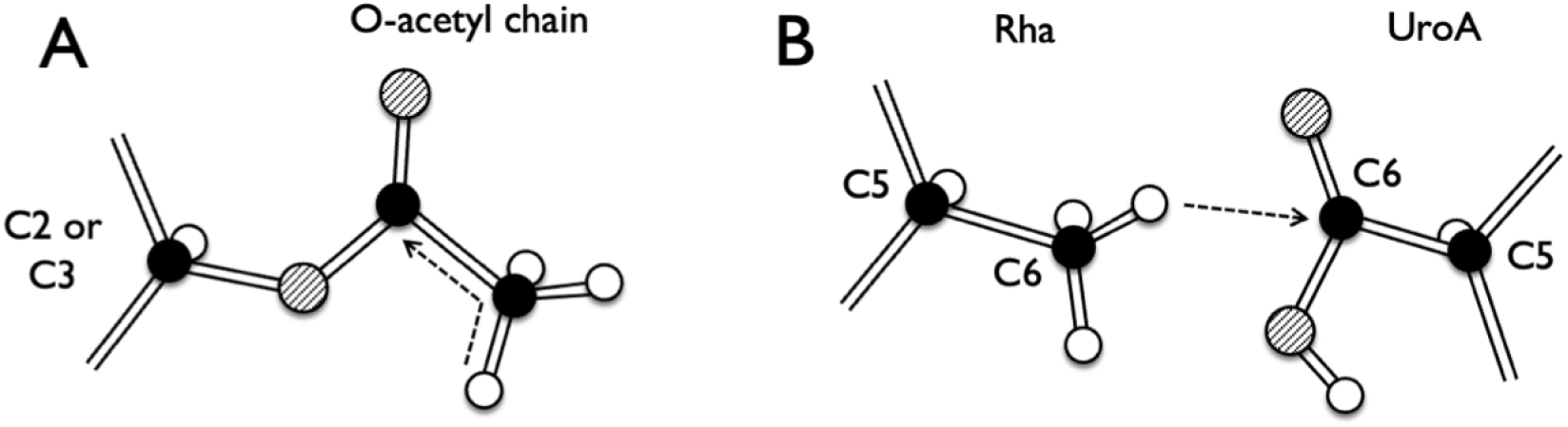
Possible magnetization transfers that would generate alkyl to carboxyl cross-peaks in a CP-HETCOR spectrum (i.e. peak #10 in Fig. 4B). (A) Through-bond transfer from H2 methyl to C1 carboxyl on *O*-acetyl sidechains, (B) Through-space transfer from H6 methyl on rhamnose residues to C6 carboxyl on uronic acids. Black circles are carbon atoms, hatched circles are oxygen atoms, white circles are hydrogen atoms; dashed arrows indicate magnetisation transfer (= “cross polarization”).

A possible alternative explanation for our observed cross-peak (peak #10 in Fig. 4B) is that it arose from through-space cross-polarization by spin diffusion, for example from rhamnose C6 -CH_3_ protons to uronic acid C6 -COOH carbons (Fig. 5B) or from cyclic pyruvyl ketal - CH_3_ protons to cyclic pyruvyl ketal -COOH carbons. However, neither of these are consistent with the ester peak seen in the FTIR spectra (Fig. 3B) and both would require a degree of alignment between extracted polymers for which there is no evidence. Additionally, we may rule out the involvement of pyruvyl ketals because spin-diffusion from rhamnose or pyruvyl ketal -CH_3_ protons to pyruvyl ketal -COOH carbons should be accompanied by concomitant spin-diffusion to the middle pyruvyl carbon (the one that cross-links the C3 and C4 O atoms of galactose). This would give a strong crosspeak in the long-contact spectrum between ^1^H 1.5-2.5ppm and ^13^C 90-100ppm, but no such peak increase was observed (Fig. 4B), which suggests that any cross-peak from cyclic ketal groups was minimal or non-existent. We also note that cyclic pyruvyl ketals have only been observed in the galactans of galactan-rich *Codium* species (Bilan et al., 2007) and that *Ulva* does not contain high levels of these polymers (Percival, 1979).

### Base-labile acetate assays suggest significant levels of acetylation in the ulvam fraction of Ulva cell walls

Both our FTIR (Fig. 3) and ^13^C NMR spectra (Fig. 4) suggest that *Ulva* cell walls contain reasonable amounts of *O*-acetylated ulvan. To quantify the extent of that acetylation, and to confirm that the acyl groups were acetyl groups, we subjected ulvan extracts (i.e. U1 and U2 fractions combined) to high temperature base hydrolysis (1M NaOH at 80ºC for 8h) and measured the liberation of acetate. Under these conditions, base hydrolysis should remove carboxylate ester-linked alkyl groups (eg *O*-acetyl groups) but would not be expected to degrade the glycosidic bonds of the polymer backbone, which would require acid hydrolysis.

As expected, base hydrolysis significantly lowered the ester peaks at ∼1730 cm^−1^ and the sulfate peaks at ∼1220 cm^−1^ (Supplemental Fig. S2) and we found that around 50-90mg of base-labile acetate was liberated from 1g of ulvan (i.e. fractions U1+U2). The molar mass of acetate is ∼59g so if we presume an average molar mass of 160-170g for *Ulva* cell wall sugar monomers, our results suggest a stoichiometry of one base-labile acetate molecule for every 4-7 ulvan monomers.

### FTIR-visible ester resonances reversibly decrease in response to metal stress, but not saline stress

To investigate the dynamic behaviour of *Ulva* cell wall *O*-acetyl modifications, we subjected *Ulva* thalli to one of three stresses: hyposaline (0‰), hypersaline (90‰) and high metal (100µM copper or 10µM lead). Each stress was imposed for 24h and then removed, leaving thalli to recover in artificial seawater (35‰) for another 24h. Thalli subjected to hyper- and hyposaline stress showed no obvious changes in their FTIR-visible ester peaks (Fig. 6B, C). However, metal-exposed thalli showed a decrease in their FTIR-visible ester peaks (Fig. 6D), which was reversed after metal-exposed thalli had been restored to control conditions for 24h (Fig. 6E).

**Figure 6.**
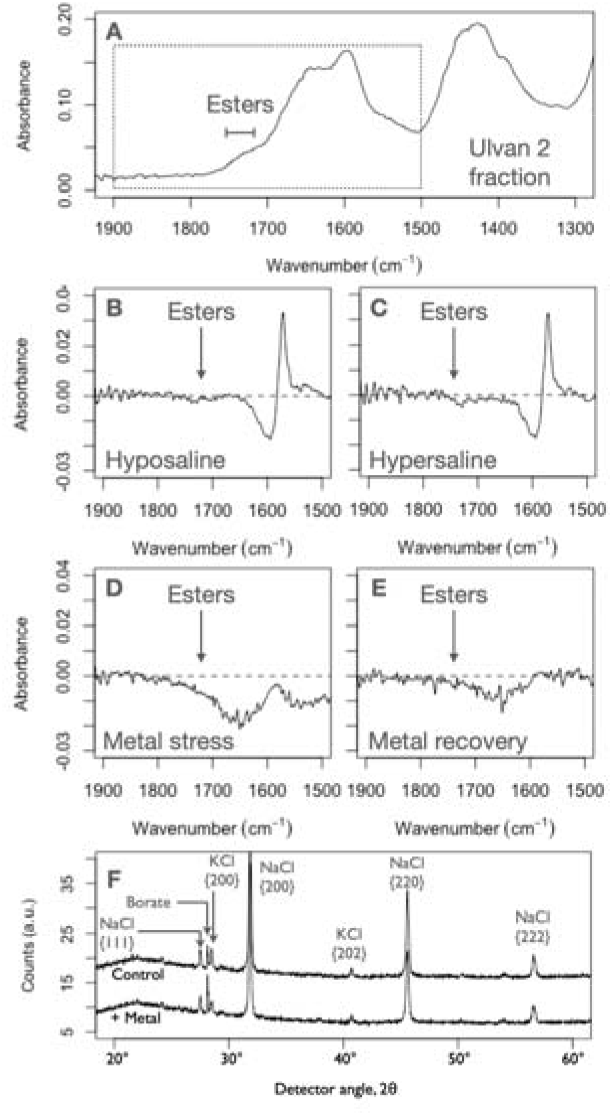
Changes in FTIR-visible ester peaks and borate levels in ulvan 2 extracts of *Ulva* thalli during stress. (A) Representative FTIR spectrum of the ulvan 2 fraction, showing the major resonances between 1900-1300 cm^−1^ and identifying the range in which ester peaks fall. The dotted box indicates the regions that are expanded in (B)-(E), (B) Representative difference spectrum generated by subtracting a paired control spectrum from the spectrum of ulvan from a hyposaline treated (0‰) *Ulva* thallus, (C) Representative difference spectrum generated by subtracting a paired control spectrum from the spectrum of ulvan from a hypersaline treated (90‰) *Ulva* thallus, (D) Representative difference spectrum generated by subtracting a control spectrum from a spectrum derived from metal treated thalli (100µM copper), showing a decrease in ester and carboxylic acid resonances after metal treatment, (E) Representative difference spectrum generated by subtracting a control spectrum from metal-recovered thalli (24h after metal removal), showing recovery of the ester signal, (F) Representative XRD scans showing an increase in borate following metal stress (100µM copper and 10µM lead give similar results).

In land plants, *O*-acetylation is found on the C2 and C3 positions of polymer sugar residues and on the C6 of Galactose and some Glucose residues in angiosperm xyloglucan. In ulvan, however, free OH groups on the rhamnose C2 and C3 positions are believed to bind significant amounts of borate after the ulvan has been exported into the cell wall (Haug, 1976). We might therefore expect reduced *O*-acetylation to free up C2 and C3 positions for borate. Accordingly, we used X-ray diffraction to look at whether the high metal-induced decrease in FTIR-visible ester signal was accompanied by complementary changes in borate. We found that high metal concentrations were accompanied by a single peak increase at a detector angle of 28.07° (Fig. 6F), which corresponds to the 3.18Å lattice distance that is characteristic of boric acid (Zachariasen, 1934). We conclude that ulvan-associated borate rises during metal stress, while ulvan *O*-acetylation falls.

### Ulva genomes contain orthologs of the Arabidopsis genes required for cell wall O-acetylation

Given our spectroscopic evidence for *O*-acetylation, we searched for orthologs to Arabidopsis Reduced Wall Acetylation, *AXY9* and Trichome Birefringence-Like genes in the sequenced genomes of the chlorophyte seaweeds *Ulva compressa, Ulva prolifera*, and *Caulerpa lentillifera*.

A single sequence that is homologous to *Arabidopsis RWA* genes exists in the genomes of *Ulva compressa* (*UM026_0056*) and *Ulva prolifera* (Fig. 7A) and has already been identified in the genome of *Chlamydomonas reinhardtii* (*CHLRE_09g389950v5*; Pauly and Ramírez, 2018). Similarly, a single sequence that is homologous to the *Arabidopsis AXY9* sequence exists in the genomes of both *U. compressa* (*UM011_0103*) and *U. prolifera* (Fig. 7B). The *Ulva* sequences cluster by phylogeny within the Cas1p family (Fig. 7A, B), so they may be regarded as orthologs. No *AXY9* orthologs exist in *Chlamydomonas* and no orthologs to either *RWA* or *AXY9* exist in *Caulerpa*, which has a cell wall built from xylan. The *Ulva compressa RWA* ortholog (*UM026_0056*) is predicted to have 12 membrane-spanning helices, while the *AXY9* ortholog (*UM011_0103*) is predicted to have one membrane spanning helix, as well as the conserved GDS and DxxH motifs (Fig. 7C).

**Figure 7.**
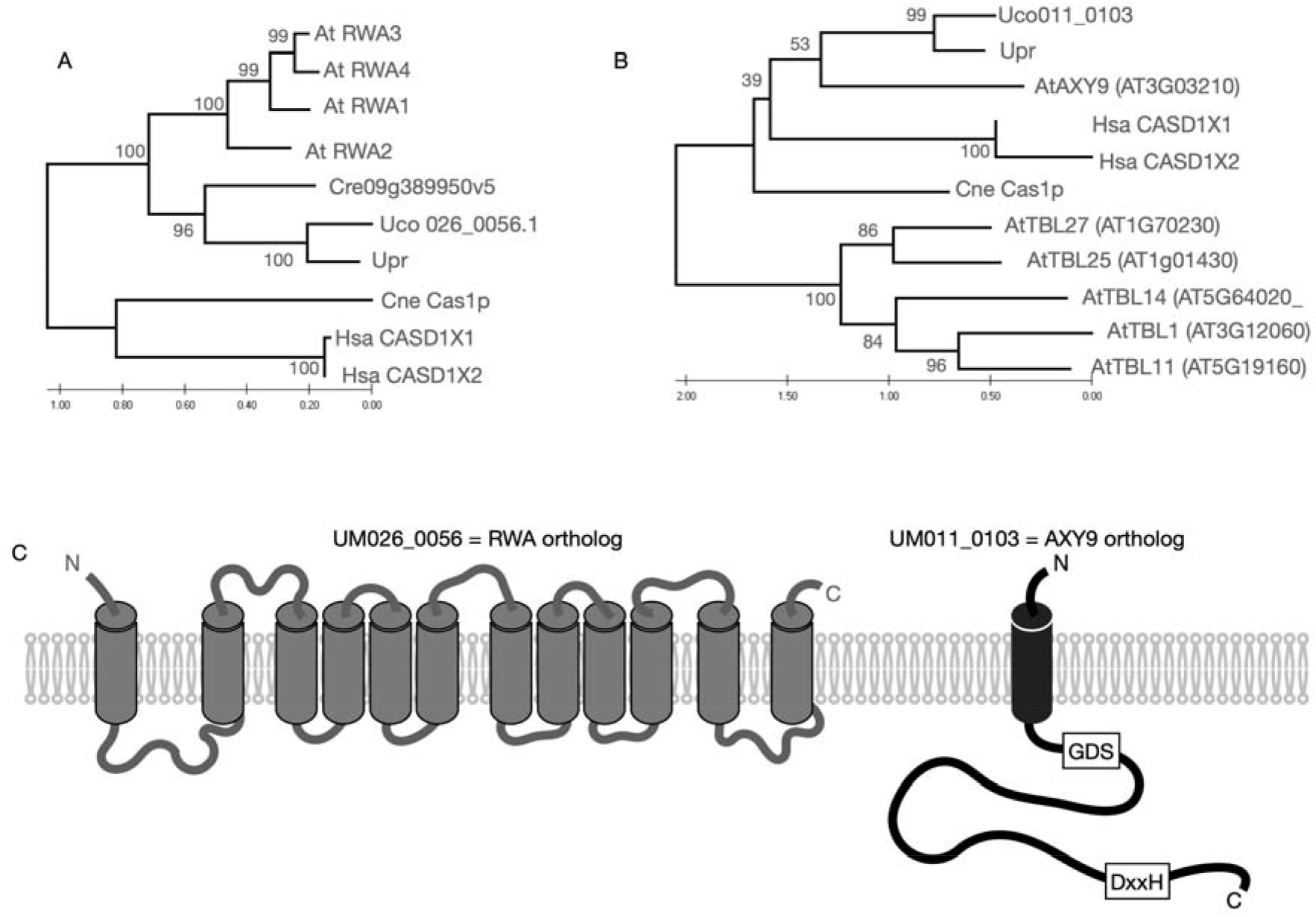
Identification of an *O*-acetyltransferase ortholog in green seaweed genomes. For and (B), trees are drawn to scale, with branch lengths measured in the number of substitutions per site. The percentage of trees in which the associated taxa clustered together is shown next to the branches. (A) Maximum Likelihood phylogenetic tree of RWA-like genes from selected species. The tree with the highest log likelihood (-8550) is shown. This analysis involved 10 amino acid sequences and a total of 798 positions in the final dataset. Maximum Likelihood phylogenetic tree of AXY9-like genes from selected species. The tree with the highest log likelihood (-9630) is shown. This analysis involved 11 amino acid sequences and a total of 740 positions in the final dataset. (C) Predicted structures of the Cas1p and TBL orthologs from *Ulva compressa*.

### The Ulva O-acetyltransferase orthologs are co-localized in the Golgi

To check whether the expressions of *U. compressa RWA* and *AXY9* orthologs were consistent with roles in golgi *O*-acetyltransferase activity, we created YFP-tagged transgenic lines for each ortholog. Both proteins localise to discrete areas close to the nucleus (Fig. 8; see also Supplemental Figs S3-S5). These localisation patterns do not correspond to those for microbodies, mitochondria, the nucleus or the ER (Blomme et al., 2021) but they are consistent with the location of the golgi in *Ulva compressa* (Løvlie and Bråten, 1968). We conclude that both the *Ulva AtRWA* ortholog, *UM026_0056*, and the *Ulva AtAXY9* ortholog, *UM011_0103*, are golgi-localised.

**Figure 8.**
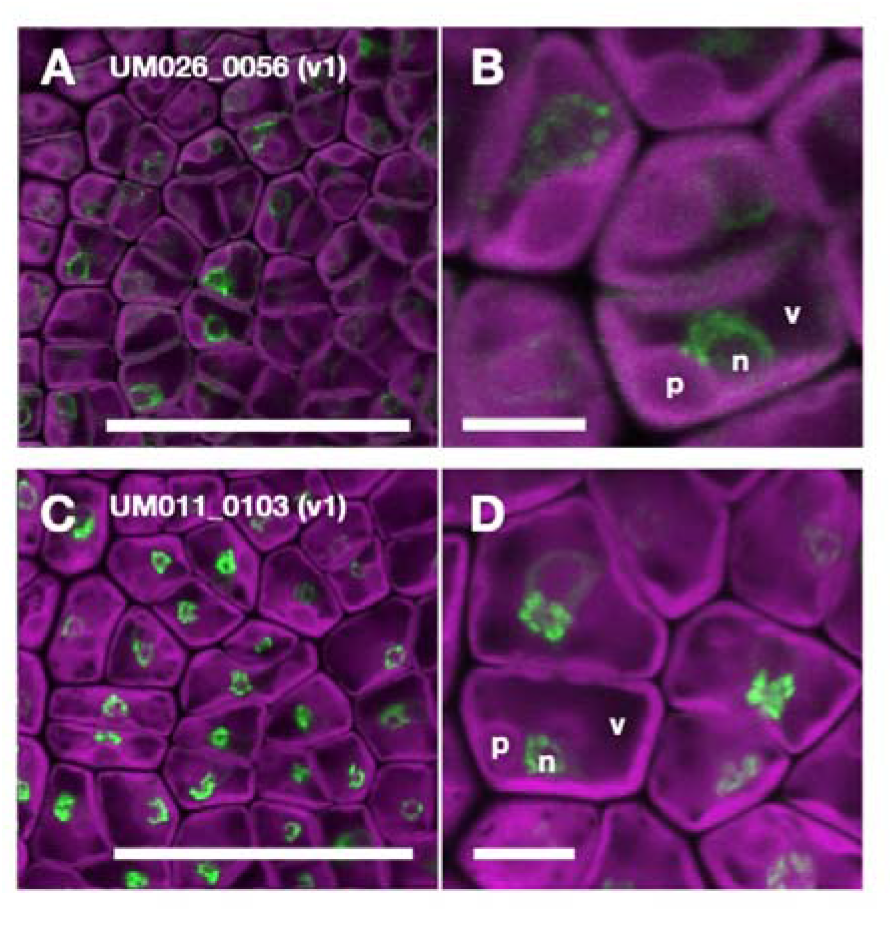
Visualization of transgenic *Ulva* lines expressing YFP-tagged orthologs to *O*-acetyltransferases. (A) and (B) YFP-tagged UM026_0056, which is the transmembrane *AtRWA* ortholog, (C) and (D) YFP-tagged UM011_0103, which is the manually predicted lumen-facing loop *AtAXY9* ortholog (see Supplemental Fig. S5). For each transgenic line, two representative individuals are shown, one at low magnification (a and c) and one at higher magnification (b and d) in which the nucleus (n), vacuole (v) and pyrenoid (p) are identified. Green indicates the YFP signal and magenta represents chlorophyll autofluorescence. Scale bar: 50μm in (A) and (C) and 5μm in (B) and (D). See also Supplemental Figs S3-S5.

### Extractable cell wall fractions show stronger FTIR-visible ester resonances in Ulva, which has acetyltransferase orthologs, than in Caulerpa, which does not

The bryopsidalean green seaweed, *Caulerpa lentillifera*, has no orthologs to *Arabidopsis* acetyltransferases. To investigate whether this is associated with a lower degree of cell wall acetylation, we compared the FTIR spectra of the slowly oxalate-extractable fractions from *Ulva* (i.e. its ulvan 2 fraction) and *Caulerpa* and found clear differences (Fig. 9). Specifically, *Ulva* has stronger peaks at ∼1730 cm^−1^ and ∼1220 cm^−1^ (Fig. 9) suggesting a greater degree of acetylation and sulfation on the ulvan 2 fraction of *Ulva*. The equivalent fraction in *Caulerpa* does, however, show a tail to the left of the ∼1620cm^−1^ carboxylic acid resonance, which suggests that the oxalate-extractable cell wall fraction in *Caulerpa* may also contain ester linkages.

**Figure 9.**
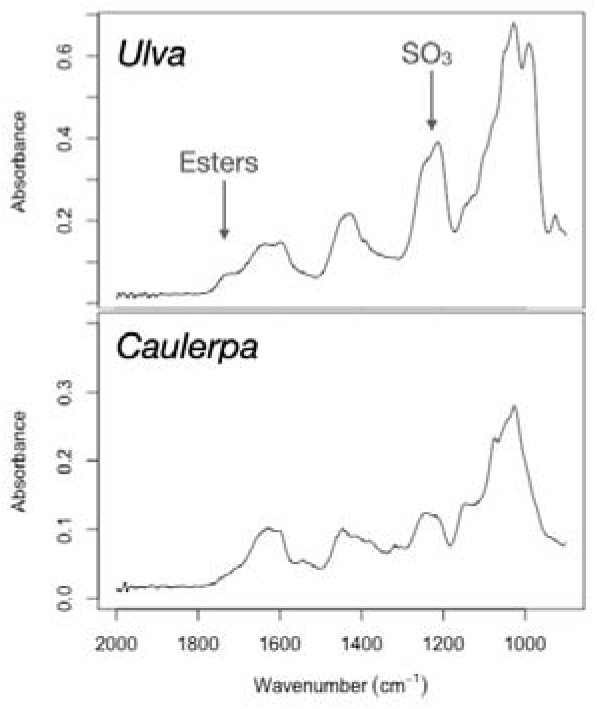
Representative FTIR spectra of the slowly oxalate-extractable cell wall fractions from the ulvalean chlorophyte seaweed *Ulva compressa* (i.e. its ulvan 2 fraction) and the bryopsidalean chlorophyte seaweed *Caulerpa lentillifera*, showing the major resonances between 2000-900 cm^−1^.

## Conclusions

Our results show that the cell walls of some chlorophyte seaweeds can be dynamically *O*-acetylated (Figs 3-6) in similar fashion to the *O*-acetylation of pectins and hemicelluloses in many land plants. We suggest that this acetylation is most noticeable on the extractable ulvan fraction (Fig. 3) and we demonstrate that the genomes of some chlorophyte seaweeds contain orthologs to known *O*-acetyltransferases (Fig. 7) that are expressed in the golgi in which the non-cellulosic cell wall fractions are made (Fig. 8). Our estimated stoichiometry of one *O*-acetyl group for every 4-7 ulvan monomers is an approximate one, but is consistent with the 30-60% levels of *O*-acetylation reported in the water extractable polymers and hemicelluloses of many land plants (Teleman et al., 2000; Capek et al., 2002; Yuan et al., 2013). This is a significant level of modification and suggests that acetylation may be almost as common a modification as the better-known 2-O-sulfate and 3-O-sulfate ester groups on xylose, rhamnose and galactose residues (Percival and Wold, 1963; Bilan et al., 2007).

The most obvious gap in our work is that gene knockout protocols have not yet been robustly developed in the green seaweeds, so we are unable to create loss-of-function transformants for either of our two candidate acetyltransferase genes. Instead, we show that a green seaweed species that has the candidate acetyltransferases (*Ulva compressa*) also has a strong ester peak, while a green seaweed species that lacks the candidate acetyltransferases (*Caulerpa lentillifera*) has a weaker or possibly non-existent FTIR-visible ester peak (Fig. 9; see also results for a red seaweed species, *Mastocarpus stellatus*, in Supplemental Fig. S6). We note that, despite its genome lacking any acetyltransferase ortholog, there is some evidence for *O*-acetylation in *Caulerpa* in the form of an ester ‘tail’ at ∼1710cm^−1^ in the FTIR spectrum of the *Caulerpa* oxalate-extractable cell wall fraction (Fig. 9). Cell wall *O*-acetylation in the absence of working *AtRWA* gene products has previously been observed in Arabidopsis knockouts (Lee et al., 2011) and our observations support the suggestion (Lee et al., 2011) that the green lineage may possess a second, as-yet-unidentified, acetylation pathway in addition to the *AtRWA/AtAXY9/AtTBL* pathway.

Our findings raise four obvious questions. First, what is the exact location of *O*-acetyl groups in ulvan? We are currently unable to answer this. Ulvan is a heterogeneous polymer and we have not yet worked out which residues the acetyl groups are attached to. In land plants, extensive 2-*O*- and 3-*O*-acetylation of sugar residues has been reported on xylans, glucomannans and pectins, with 6-*O*-acetylation on xyloglucans (Gille & Pauly, 2012), but land plants have the 50 or so members of the *TBL* gene family, which may allow them to target a broader range of sugars than *Ulva*, which has only the one *AXY9* ortholog. Nonetheless, the AXY9 protein is thought to be promiscuous, so we expect that acetylation in *Ulva* will occur on a range of sugar residues.

Second, and given the possible promiscuity of the *AXY9* ortholog, is acetylation truly confined to ulvan? The FTIR-visible ester signal in *Ulva* is only visible on the slowly oxalate-extractable ulvan 2 fraction (Fig. 3) but the hemicellulose b fraction was prepared through base hydrolysis and strong base hydrolysis will remove acetic esters. The lack of any apparent ester signal in the hemicellulose b fraction (Fig. 3C) may, therefore, reflect the removal of ester groups during hemicellulose extraction rather than any actual lack of acetylation. To set against that, there are no strong FTIR-visible ester resonances in the ‘not oxalate-extractable’ residue (Fig. 1), which was not subjected to base hydrolysis (Supplemental Fig. S7). Taken together, we cannot rule out the possibility of some *O*-acetylation on *Ulva* hemicelluloses but our evidence suggests that acetylation is predominantly on the extractable ulvan polymers.

Third, what is the role of cell wall *O*-acetylation in *Ulva*? Again, we cannot currently answer this question. *O*-Acetylation is known to change the rheological properties of polysaccharides (Cardoso et al., 2003) and the decrease in acetylation seen during *Ulva* metal stress (Fig. 6) is similar to that seen in land plants in response to aluminum stress (Zhu et al., 2014), but the functional significance of many algal cell wall modifications remains a matter of some conjecture (Domozych et al., 2012). We would like, however, to draw attention to the possible interactions between acetylation and the addition or removal of the borate and sulfate sidechains in ulvan (Fig. 6F). Ulvan that is rapidly oxalate-extractable (i.e. the ulvan 1 fraction) is less acetylated but more sulfated than ulvan that is slowly oxalate-extractable (Fig. 3A, B) and a decrease in ulvan acetylation during metal stress is mirrored by a rise in borate levels.

Fourth, what light do our findings shed on the evolution of cell wall *O*-acetylation? We can find no obvious orthologs to the *AtRWA* or *AtAXY9* genes in the genomes of the red seaweeds *Chondrus crispus* and *Porphyra umbilicalis* and we found no strong evidence for FTIR-visible ester resonances in any of the cell wall fractions of the red seaweed *Mastocarpus stellatus* (Supplemental Fig. S6), which is closely related to *Chondrus crispus* (both are members of the same order, the Gigartinales). We therefore tentatively presume that the presence of *AtRWA* and *AtAXY9* orthologs in the *Ulva* genome means that this system evolved before the divergence of the chlorophyte and streptophyte lineages perhaps 800 MYA (Kumar et al., 2017) but why has acetylation then been lost by *Caulerpa* and why has *Chlamydomonas* retained its *AtRWA* ortholog?

Finally, we note one caveat to our current work: we cannot rule out the additional involvement of longer alkyl chains, possibly propionyl or butyryl chains, in *Ulva* cell wall modification. Our CP-HETCOR spectra are complex and contain small uncharacterized resonances that would be consistent with the presence of longer alkyl chains (eg the small shoulder peak around 26ppm in Fig. 4); the acetate kinase used in our assays will phosphorylate propionate, albeit much more slowly than it phosphorylates acetate (Winzer et al., 1997), propionylation is known to occur on histones and to be catalysed by the same enzymes as acetylation (Liu et al., 2009), and the exact substrate specificity of the Cas1p family of *O*-acetyltransferases remains unclear.

## Methods and Materials

### Seaweed collection and cultures

For cell wall extractions, *Ulva* and *Mastocarpus* thalli were collected from the foreshore of Seaham beach, County Durham, UK, with *Ulva* being provisionally identified as *Ulva compressa* because of its location, ribbon-like morphology and hood-shaped plastids (Brodie et al., 2007). *Caulerpa lentillifera* was ordered from Planted Reef, Taunton, UK. Seaweeds were maintained in artificial seawater (Sigma Aldrich, Darmstadt, Germany) in 250mL Erlenmeyer flasks at 8ºC under a 16h:8h light:dark cycle.

Transgenic lines were generated using established cultures of *Ulva mutabilis* Føyn “slender” mutant strain (Føyn, 1958), which is conspecific with *Ulva compressa* (Steinhagen et al., 2019). Individuals were cultivated under long day conditions (16h light:8h dark; 21°C; 75µM; Spectralux Plus NL-T8 36W/840/G13 fluorescent lamp) in standard Petri dishes containing synthetic Ulva Culture Medium (UCM; Stratmann, Paputsoglu and Oertel, 1996; Boesger, Kwantes and Wichard, 2018). *Ulva* was grown and parthenogenetically propagated as described previously (Wichard and Oertel, 2010).

### Extraction and separation of cell wall fractions

Total cell wall extraction was carried out using standard alcohol-insoluble residue methods: thalli were frozen in liquid nitrogen and ground to a fine powder, still under liquid nitrogen. The powdered sample was washed twice to remove pigments, in 96% and 100% ethanol respectively; after each wash the sample was centrifuged for 15min at 3,000g and the supernatant was discarded. The remaining pellet was washed twice more in 2:3 methanol:chloroform solution to remove polar and nonpolar metabolites; each wash ran overnight with shaking, and was followed by 15min centrifugation at 3,000g with the supernatant being discarded. Finally, the pellet was washed three more times in 100%, 65%, and 80% ethanol solutions; after each wash the suspension was centrifuged for 15min and the supernatant was discarded. The remaining pellet was resuspended in 100% ethanol and dried to leave cell wall powder.

Cell wall fractions were then extracted and separated from cell wall powder using a modification of the method of O’Rourke and colleagues (2015; Fig. 1). The ‘ulvan’ fractions were solubilized with 0.2 M ammonium oxalate (pH 4.0–4.3) at 100°C for 2 h with the supernatant being collected as ‘ulvan 1’ or U1, followed by a further 16 h with the supernatant being collected as ‘ulvan 2’ or U2 (Fig. 1). These rapidly and slowly oxalate extractable extracts (U1 and U2) were dialysed and freeze-dried. The precipitated residue that remained in the pellet after oxalate extraction was washed 3x in water to remove oxalate buffer and kept as the ‘not oxalate-extractable’ material (Fig. 1).

In some cases, hemicelluloses were solubilised from the ‘not oxalate-extractable’ residue by extraction in 6 M NaOH for 72 h at 37°C (Edelmann and Fry, 1992). The supernatant was neutralized with acetic acid and dialysed against water; material that precipitated during these operations (termed hemicellulose a; Ha) was sedimented by centrifugation at 5000 g for 10 min, rinsed in water and freeze-dried. The remaining solution (hemicellulose b; Hb) was also freeze-dried. The material that was non-extractable in NaOH material was rinsed several times with pH 4 buffer, and the pooled washings (wash ‘W’) were dialysed and freeze-dried. The final residue (‘α-cellulose’, αC) was rinsed in water and freeze-dried.

### Polysaccharide hydrolysis and chromatography of sugars

Cell wall fractions (5mg) were hydrolysed with 1mL of 2M trifluoroacetic acid at 120°C for 1h. The hydrolysate was dried, re-dissolved in water and chromatographed. Thin-layer chromatography was on Merck silica-gel plates, usually in butan-1-ol/acetric acid/water (BAW; 4:1:1) or ethyl acetate/pyridine/acetic acid/water (EPAW; 6:3:1:1); sugars were stained with thymol/sulfuric acid (Jork et al., 1994).

### Fourier-transform infrared (FTIR) spectroscopy of cell wall fractions

Fourier-transform infrared spectroscopy was carried out on a PerkinElmer Frontier IR/NIR spectrometer equipped with UATR polarization and operating with a gauge force of 60N.

### Polymer characterisation using cross-polarisation heteronuclear correlation NMR (CP-HETCOR)

Carbon-13 cross-polarization heteronuclear correlation NMR spectroscopy with Magic Angle Spinning was carried out at 100.56MHz using a 9.4T Varian VNMRS spectrometer at ambient temperature (25°C) with a 4mm PENCIL rotor spinning at 10-14kHz, contact times of 0.1 and 1.0ms, and a recycle delay of 1s. Over 1,000 transients were acquired for each spectrum and the TPPM pulse sequence was used for proton decoupling. The CP-HETCOR experiments involved Lee-Goldburg operation, with 48 t1 points spanning up to 3.49ms. Separate HETCOR experiments used contact times of 0.1ms (for detecting directly-bonded CH groups) and 1.0ms (for two-bond groups). Calibration of the proton shift scale for the HETCOR plots is technically demanding and was done before and after the HETCOR experiments, with results being transferred to spectra via the HETCOR spectra of the mixture. A sample of glycine was used for calibration purposes, with its known proton shifts of 3.0, 4.2 and 8.4ppm.

### Base hydrolysis and labile acetate assays

We used overnight base hydrolysis to remove base-labile acetyl groups from oxalate-extracted cell wall fractions. 50mg of freeze-dried fractions were resuspended in 1mL of 1M NaOH before being placed in a shaking water bath for 8h at 80ºC. After the first hydrolysis the samples were centrifuged for 15min at 3,000g and the supernatant was collected and neutralized to pH ∼7.0. The pellet was resuspended in 1mL of 1M NaOH and a second hydrolysis was performed, with the supernatant again being collected and neutralised to Ph ∼7.0. The amount of acetate in the supernatants was then measured with the K-ACETAK assay kit, according to the manufacturer’s instructions (Megazyme.com); this kit uses coupled acetate kinase/pyruvate kinase/lactate dehydrogenase reactions in a quantitative colorimetric assay.

### Imposing saline and metal stresses

The artificial seawater (Sigma Aldrich, Darmstadt, Germany) in which seaweeds were maintained has a salinity of 35‰. For hyposaline stress (0‰), thalli were transferred to pure water, for hypersaline stress (90‰) they were transferred to artificial seawater supplemented with NaCl, because seasalts on their own will not dissolve fully at high concentrations. Metal stress was induced by transferring thalli to artificial seawater supplemented with either copper chloride or lead acetate (both from 10 mM stock solutions).

### X-Ray Diffraction (XRD) analysis of borate levels

For XRD, finely ground samples were analysed using the Cu-Kα source (0.154 nm) of a Bruker AXS D8 Advance X-ray diffractometer. Peaks were identified by comparison to known standards after converting the 2θ detector angle into lattice distances using Bragg’s equation.

### Phylogenetic analysis

Reciprocal BLAST searches were conducted for homologs to the four Arabidopsis *Reduced Wall Acetylation* protein sequences (NP_568662.1, NP_001118592.1, NP_180988.3, NP_001031111.1) and the Arabidopsis AXY9 sequence (NP_186971.1), looking in the genomes of the green chlorophytes *Chlamydomonas reinhardtii* (Merchant et al., 2007), *Caulerpa lentillifera* (Arimoto et al., 2019), *Ulva prolifera* (He et al., 2021) and *Ulva compressa* (De Clerck et al., 2018) and those of the red seaweeds *Chondrus crispus* (Collén et al., 2013) and *Porphyra umbilicalis* (Brawley et al., 2017). Multiple alignments were generated in Jalview using full-length sequences in Tcoffee (Waterhouse et al., 2009). The evolutionary history was inferred using the Maximum Likelihood method and JTT matrix-based model (Jones et al., 1992), running in MEGA X (Kumar et al., 2018). Initial trees for the heuristic search were obtained automatically by applying Neighbour-Join and BioNJ algorithms to a matrix of pairwise distances estimated using the JTT model and then selectibg the topology with superior log likielihood value. Proteins included were *Cryptococcus neoformans* Cas1 (NP_568628.1), *Homo sapiens* CASD1 (N-acetylneuraminate 9-*O*-acetyltransferase) isoforms 1 (XP_011514797.1) and 2 (NP_001350355.1). Proteins included in the *AXY9* gene family tree were the Trichome Birefringence Like family genes AT3G12060 TBL1 (NP_187813.1), AT1G70230 TBL27 (NP_177180.1), AT1G01430 TBL25 (OAP18729.1), AT5G64020 TBL14 (NP_201207) and AT5G19160 TBL11 (NP_197417.1). Proteins included in the *RWA* gene family tree were AT5G46340 RWA1 (NP_568662.1), AT3G06550 RWA2 (NP_001118592.1), AT2G34410 RWA3 (NP_180988.3), AT1G29890 RWA4 (NP_001031111.1), all from *Arabidopsis*, and CnCas1 (XP_568628.1) from *C. neoformans*. All ambiguous positions were removed for each sequence pair (pairwise deletion option). Transmembrane domains were predicted using InterProScan (Finn et al., 2017).

### Ulva *transformation*

*Ulva* gametes were transformed using the PEG-mediated protocol described by Oertel, Wichard and Weissgerber (2015) and Boesger, Kwantes and Wichard (2018). For each transformation, 5µg of plasmid DNA was used per construct. Plasmids were purified using a GeneJet Plasmid Miniprep kit (Thermo Scientific). 50µg/mL Phleomycin (Invivogen) was added 48h after transformation; resistant germlings develop parthenogenetically from the transformed gametes and can be detected after 2-3 weeks of cultivation. For each transformation experiment, non-vector controls were included to confirm proper selection.

### Vector assembly

Vector assembly is based on the GreenGate cloning system (Lampropoulos et al., 2013). Sequences for entry modules were obtained via PCR-amplification of the target DNA sequence using primers that contain a ∼20bp overlap with the entry module using high-fidelity polymerase Q5 (NEB). For *UM011_0103*, two potential isoforms were cloned with only the manually curated isoform showing expression (Supplemental Figs S4, S5). After gel electrophoresis, the amplicon is purified (Zymoclean Gel DNA Recovery Kit, Zymo Research) and mixed with the pre-digested (BsaI, Thermo Scientific) entry clone and NEBuilder (NEB). After assembly (15 minutes at 50°C), the mix is transformed into chemically-competent DH5α *Escherichia coli* cells and plated out on LB+100µg/mL Carbenicillin. PCR-mediated silent mutation was introduced to remove an internal BsaI recognition site in *UM026_0056*. All *Ulva* gene sequences were cloned from genomic DNA, isolated via Omniprep for Plant (G-Biosciences).

For a Golden Gate reaction, 100ng of each entry and destination vector are assembled in one reaction mix containing 10U BsaI-HF-v2 (NEB), 200U T4 Ligase (NEB), 1mM ATP (Thermo Scientific) and 1x Cutsmart buffer (NEB). The reaction mix is incubated at 30 x (3 min @18°C and 3min @37°C) followed by 5 minutes @50°C and 5 minutes @80°C. After assembly, the mix is transformed in DH5α *E. coli* cells and plated out on LB+25µg/mL Kanamycin. Vectors were validated using Sanger sequencing (Mix2seq, Eurofins) and restriction digest using NcoI (Promega) and PstI (Promega).

All entry clones in this study are designed as described by Lampropoulos and colleagues (2013). The destination vector used is a high-copy number pGG-pUM140_0016-BE-NOST-PIBT3 containing a ccdB/CmR cassette, generated earlier (Blomme et al., 2021). All vectors generated in this study (Table S1) are available at https://gatewayvectors.vib.be. All primers used in this study are listed in Table S2.

### Confocal imaging of YFP-tagged transgenic Ulva lines

Individual thalli of one-month-old resistant individuals were selected for imaging. Detailed imaging of tissues and cells was performed using Olympus Fluoview (FV1000) confocal microscope with Fluoview software. The following excitation and emission settings (nm) were used: YFP, 515 excitation with 530-545 emission and chlorophyll, 559 excitation with 650-750 emission. For each transgenic line, at least eight independent individuals were screened, and a representative image is selected for all lines with a functional construct.

## Acknowledgements

JHB thanks the BBSRC for financial support through the SuBBSea grant (BBK020552). Studentship support came from BBSRC (AJG), the EPSRC SOFI CDT (MR), CONACYT (ATL) and the GCRF (AA). JB thanks Thomas B. Jacobs and Olivier De Clerck for facilitating the generation of tagged lines. During the 2020-21 COVID pandemic NMR, FTIR and XRD spectra were acquired by David C. Apperley, Lenny W. Lauchlan and Gary Oswald, respectively (Dept of Chemistry, Durham University). We are extremely grateful for their technical skill and support.

## Author contributions

JHB designed the research and analyzed data; AJG, MR, PJB, ATL, AA, SCF, GC and JB performed research and analyzed data. All authors contributed to the writing.

